# The Old Genetically Heterogeneous Mouse Model Recapitulates Chronic and Persistent Idiopathic Pulmonary Fibrosis with Strong Senescence Signatures

**DOI:** 10.1101/2025.07.22.665978

**Authors:** Apoorva Shankar, Gabriel Meca-Laguna, Anna Barkovskaya, Michael J Rae, Amit Sharma

## Abstract

Idiopathic Pulmonary Fibrosis (IPF) is a chronic and progressive lung disease that primarily afflicts people over the age of 65. IPF is characterized by lung scarring, an elevated senescence burden, and interstitial pneumonia, resulting in disability and mortality. The growing aging population worldwide, limited effectiveness of current treatments, and high economic burden underscore the need for robust models to investigate the underlying mechanisms and test novel interventions. Despite broad preclinical use, the bleomycin-induced murine model has notable limitations. Animals subjected to a single dose of bleomycin administration undergo recovery in weight and behavior between 21 and 28 days after administration, which is contrary to the progressive nature of IPF. Previous reports have shown that repetitive instillation of bleomycin phenocopies this aspect of the human disease. However, these methods are time-consuming and complex. Although IPF is typically associated with advanced age, most research is conducted in 6 to 8-week-old mice, which lack the age-related structural and metabolic deficits seen in humans. In this study, we report an improved model of IPF using 17-month-old UM-HET3 mice subjected to a single oropharyngeal bleomycin dosing that better mimics the persistent nature of the disease. Lung histology and immunohistochemistry (IHC) confirm persistence of fibrosis and senescence in mice 10 weeks after bleomycin administration. Furthermore, bulk RNA sequencing (RNA-Seq) analysis revealed a distinct set of gene expression signatures that is more consistent with chronic human fibrosis. This model offers greater insight into IPF pathogenesis, and we anticipate that it will enhance confidence in the human translatability of candidate therapeutic interventions.

## Introduction

Idiopathic Pulmonary Fibrosis (IPF) is a chronic and progressive interstitial lung disease with poor prognosis and a high mortality rate (Zheng et al., 2022; Hutchinson et al., 2014). It is characterized by irreversible loss of lung function due to fibrosis. The symptoms range from cough and dyspnea that worsen over time to severe limitations on physical function and independent living (Luppi et al., 2021). IPF is a disease of aging, with most individuals being diagnosed after the fifth decade of life and a higher incidence in the sixth and seventh (Selman et al., 2016). As with other diseases of aging (Rae et al., 2010), the number of people living with IPF has increased in recent decades, coinciding with global demographic aging, and national (USA) prevalence has risen by approximately 6.2% per year, on average, with each year of age (Maher et al., 2021). The crude overall incidence rate of IPF is 13.2–26.1 per 100,000 person-years (depending on the definition used) (Kondoh et al., 2022). Reports show higher occurrence in males than in females (Sankari et al., 2025; Ruscitti et al., 2020; Mulugeta et al., 2015).

The median survival for patients with IPF is typically between 3 and 5 years after diagnosis. Treatment with antifibrotic medications, such as pirfenidone or nintedanib, can delay the progression of the disease and potentially improve survival, extending it to 7 and 8 years (Ye et al., 2023; Henderson et al., 2020; Fujimoto et al., 2016).

IPF is primarily characterized by the injury of alveolar epithelial type II cells (AT2), the accumulation of inflammatory cells, and the development of fibrosis (scarring) in the lung tissue (Zhang & Wang., 2023; Henderson et al., 2020; Moore et al., 2013). The interaction between inflammatory cells, whether resident or recruited, and structural cells leads to an increase in myofibroblasts, persistent inflammatory infiltrates, and abnormal collagen accumulation (Moore et al., 2013; Sime et al., 2001; Adamson et al., 1998; Coultas et al., 1994). More recently, cellular senescence has been causally linked to IPF. Senescent cells secrete senescence associated secretory phenotype (SASP) factors and impair epithelial repair, leading to a more chronic and irreversible nature of the disease (Yamada et al., 2022; Zhang & Wang., 2023; Yao et al., 2021). Markers of cellular senescence are detected in IPF lung tissue, and removal of these senescent cells is shown to improve lung health in mice (Liu RM & Liu G., 2020; Schafer et al., 2017). Importantly, pilot studies report that treatment with senolytics in human patients improves clinical symptoms (Justice et al., 2019).

The rising prevalence, high levels of morbidity and mortality, and limited therapeutic options for IPF motivate research to increase the understanding of the pathogenesis and underlying mechanisms of the disease and to develop novel therapies. In attempts to do so, researchers have heavily relied on animal models (Moeller et al., 2007). Among these, the most common involve administration of a variety of noxious substances, including silica, fluorescein isothiocyanate (FITC), and radiation. The best-characterized model involves the intratracheal or intranasal delivery of a single dose of bleomycin (Gul et al., 2023; Limjunyawong et al., 2014; Degryse et al., 2011).

In addition to models based on the administration of toxic substances, models that involve the overexpression of cytokines such as TGF-β, TGF-α, IL-13, IL-1β, and TNF-α have been explored. However, these models do not reflect the endogenous regulation seen in humans (Tashiro et al., 2017). Additionally, humanized models of IPF involving immunodeficient non-obese diabetic/severe combined immunodeficient (NOD/SCID) mice have enabled the engraftment of human IPF lung cells into these mice, resulting in the progression of fibrosis. Despite this, the lack of a functional adaptive immune response in these mice limits their relevance and does not accurately reflect the immunological context in humans (Moore et al., 2017; Pierce et al., 2007). Overall, these models are time-intensive, expensive, and complex, limiting their accessibility across research laboratories.

Bleomycin, a complex glycopeptide primarily isolated from the actinobacterium *Streptomyces verticillioides,* has been widely adopted as an antitumor antibiotic to treat various carcinomas and lymphomas. It is often administered to develop a murine model of IPF. Bleomycin disrupts DNA synthesis and replication by causing single and double strand breaks in the DNA, leading to cell cycle arrest in tumor cells. This process occurs through the chelation of metal ions, followed by a reaction between the resulting pseudoenzyme and oxygen, which leads to the formation of superoxide and hydroxide free radicals that cleave DNA (Claussen & Long, 1999).

The absence of bleomycin hydrolase, an enzyme that inactivates bleomycin, in the skin, lungs, and mucous membranes leads to bleomycin-induced toxicity in these organs (Liu et al., 2017; Umezawa et al., 1966). Excessive production of reactive oxygen species can trigger an inflammatory response, leading to pulmonary toxicity, the activation of fibroblasts, and ultimately, the development of fibrosis (Chaudhary et al., 2006; Hay et al., 1991). These properties make it a valuable basis for an animal model of IPF.

Bleomycin induces two main phases shortly after drug administration: an inflammatory phase and a fibrotic phase. Studies have reported an early upregulation of pro-inflammatory cytokines, such as tumor necrosis factor-α, interleukin-1, interleukin-6, and interferon-γ, followed by the expression of pro-fibrotic markers, including procollagen-1, transforming growth factor-β1, and fibronectin, which peaks around day 14. The transition between the two phases occurs approximately 7-10 days post-administration (Barravecchia et al., 2024; Moeller et al., 2008; Chaudhary et al., 2006).

None of the currently available models of IPF fully recapitulate all aspects of the human disease or the classical histopathology of IPF, thereby limiting their translational potential (Moore et al., 2013; Moore et al., 2008; Moeller et al., 2007). Among the limitations of the bleomycin model, mouse strains vary in their susceptibility to the bleomycin-induced fibrosis: C57BL/6J and CBA are strong responders, while BALB/cJ mice are comparatively resistant (Phan & Kunkel., 1992). This strain-specificity limits the replicability of findings in other mouse strains. Another significant limitation of this method is that the young (6–8-week-old) mice used in the standard protocol show signs of recovery from a single instillation of bleomycin, with fibrosis and disease symptoms often returning to nearly normal levels within 3-4 weeks, making it ill-suited to model a progressive chronic disease. Moreover, young animals have not undergone the multi-system, complex effects of the aging process to which human IPF patients are subject (Collard HR, 2010), which likely means that they are deficient in many of the contributors to the clinical condition of patients and the intractability of the human disease. Additionally, this capacity for spontaneous resolution makes it challenging to distinguish between the effects of the drug and the natural repair process of the lung (Redente et al., 2021; Chung et al., 2003; Izbicki et al., 2002). Some studies have reported that repetitive small doses of bleomycin (instead of the standard single instillation protocol) induce more severe fibrosis that persists for 3-6 months (Limjunyawong et al., 2014; Redente et al., 2021; Degryse et al., 2010). However, these protocols are both time-consuming and complex.

These limitations led us to explore the possible advantages of an IPF model based on the administration of bleomycin to aged mice, as IPF is often described as a disease of the elderly. Older animals are reported to have higher susceptibility to bleomycin-induced injury compared to younger mice and are shown to mimic the progressive nature of the disease (Stout-Delgado et al., 2016; Sueblinvong et al., 2012; Redente et al., 2011). Additionally, aged male mice are reportedly more susceptible to bleomycin-induced pulmonary fibrosis than females, which mirrors the sex disparity of human IPF (Redente et al., 2011).

To address the risk that findings made in a particular strain of mice may not be generalizable due to their long-inbred, often uniformly homozygous genetics, we opted to use a genetically diverse four-way cross mouse model known as UM-HET3 instead of an established strain of mice in our model. UM-HET3 mice are the second-generation offspring of (BALB/cJ × C57BL/6J) female progenitors and (C3H/HeJ × DBA/2J) male progenitors, leading to diverse but reproducible genetic backgrounds and variability in age-related mortality (Flurkey et al., 2010). The National Institute on Aging (NIA) uses UM-HET3 mice in its Interventions Testing Program (ITP) (Nadon et al., 2008). Lifespan studies and testing of various anti-aging interventions are conducted across three independent sites to enhance the reliability and robustness of the results. Drugs such as aspirin and rapamycin have been reported to show significant lifespan extension among others (Nadon et al., 2017; Lamming et al., 2013; Miller et al., 2011). It is likely that this genetic diversity better reflects human populations, with variable susceptibility to age-related diseases across the population.

In this study, we demonstrated that a single dose of bleomycin administered via the oropharyngeal route to old (17–18-month-old) UM-HET3 mice leads to a chronic and persistent model of IPF, reflecting key aspects of the human disease. We utilize immunohistochemistry, gene expression analysis, the open field test, and bulk RNA sequencing (RNA-Seq) to characterize this mouse model.

## Materials and Methods

### Animals and the bleomycin model

The use of animals was reviewed and approved by the Institutional Animal Care and Use Committee (IACUC) protocol (SRF-01.8). All procedures used in this study adhered to the ethical guidelines for animal research. Male UM-HET3 mice (16-17-month-old) weighing 38-57 g were used for this study. All animals were maintained in a 12-hour light/dark cycle with standard chow and water available ad libitum. A total of 37 mice were used in this study. They were divided into two groups: a control group and a treatment group. A single dose of bleomycin (MedChem Express, cat no. HY-17565) resuspended in saline (BioWorld, cat no. 40120975-2) was administered to anesthetized mice (50 µL) in the treatment group using the oropharyngeal route of administration at 2.5 (n=11) or 3.5 mg/kg of mouse body weight (n=17). Mice in the control group (n = 9) received an equal volume of saline solution.

### Open field test

The open field test has previously been identified as a valuable tool for assessing anxiety-like behavior and reduced locomotor activity in bleomycin-induced IPF models. Mice treated with bleomycin spend significantly less time in the center and move less frequently, possibly due to systemic inflammation and injury (Xiong et al., 2023). Open field data were collected for sham- and bleomycin-treated mice at baseline, day 15 (D15), and day 23 for group 1 (D24). Similarly, baseline, D15, D23, D46, and D71after bleomycin administration for group 2 (D72). Mice were acclimated in the testing room for a minimum of 1 hour before measurements were taken. Noldus Ethovision XT was used to measure the animal’s movement and spontaneous activity in a controlled environment. The mice were placed in the field of view (also known as the arena) and allowed to explore for 10 minutes. The total distance was recorded and calculated using the Ethovision software, which was later used for statistical analysis.

### Tissue analysis

Animals were randomly assigned to two groups for tissue collection. Lung tissue was collected from group 1 on day 24 after bleomycin instillation and the remaining animals (group 2) on day 72. For validation of fibrosis, the left lung was collected for histology, and the flash-frozen right lung was used for quantitative reverse transcription PCR (RT-qPCR). The study excludes mice that lost less than 5% of their body weight in the first 14 days and euthanized mice that have reached humane endpoints (Colville et al., 2023).

### Histology

Lung samples were collected at both time points after bleomycin administration; the left lung was fixed in 10% formalin buffer (Sigma-Aldrich, cat no. HT501128) for 24 hours for histological analysis. The right lung was flash frozen in liquid nitrogen and stored at -80 °C for further use. Paraffin embedding of the left lung tissue was done at Zyagen (San Diego, USA). Picrosirius red staining was performed to evaluate the overall morphology and assess fibrosis in the lungs. The paraffin tissue blocks were sectioned at a thickness of 5 µm, deparaffinized, rehydrated, and treated with a 0.2% phosphomolybdic acid aqueous solution for 5 minutes. The sections were stained with Picrosirius red stain in 0.1% saturated aqueous solution of picric acid for 2 hours at room temperature. The sections were washed with 0.015 N hydrochloric acid for 3 minutes, rinsed in 70% ethanol, and then dehydrated with increasing concentrations of ethanol and xylene for 2 minutes and 5 minutes, respectively. Sections were mounted and allowed to dry before imaging. The sections were imaged using an Olympus SlideView VS200 and quantified using ImageJ. Lung tissue from sham-treated mice was pooled during analysis from both time points.

Hematoxylin and Eosin (H&E) staining was performed using the Vector Laboratories (cat no. H-3502) staining kit. Sections were deparaffinized and rehydrated with xylene and ethanol. The staining was performed as per the manufacturer’s instructions.

### Immunohistochemistry

Mouse left lungs were fixed in formalin (10%) and embedded in paraffin at the Zyagen facility (San Diego, USA). Formalin-fixed paraffin-embedded (FFPE) tissue blocks were sectioned into 5 µm-thick sections. The slide containing the sections was deparaffinized using xylene and then rehydrated through a graded ethanol series. Antigen retrieval was performed by heating (1 minute in the microwave) the buffer in sodium citrate buffer (pH 6.0) and immersing the slides for 10 minutes. The endogenous peroxidase activity was quenched using 3% hydrogen peroxide (Sigma-Aldrich, cat no. 216763-500ML) in deionized water. Following, Tris-buffered saline with 0.1% Tween 20 (1x TBST, Bio-Rad cat no. 1706435) was used as a wash buffer. To reduce nonspecific binding, samples were blocked with 5% goat serum in TBST for 1 hour at room temperature. The tissues were incubated overnight at 4 °C with a 1:50 dilution of primary p21^WAF1/Cip1^ antibody (Thermo Fisher, cat no. 14-6715-81). HRP-conjugated goat anti-mouse secondary antibody (Vector Laboratories, cat no. MP-7451-15) was added for 30 minutes at room temperature. Detection was carried out using a DAB substrate kit (Vector Laboratories, ImmPACT DAB Substrate Kit, cat. no. SK-4105) until the desired intensity was achieved. The sections were then counterstained with hematoxylin (Vector Laboratories, cat no. H-3401-500) and imaged using the Olympus SlideView VS200 Scanner at 20x magnification. Four different fields of view were obtained per sample for quantification.

The same protocol was followed for staining tissue sections with SPP1 (Proteintech, cat no. 22952-1-AP) at a 1:300 dilution overnight at 4°C. Sham samples were pooled during analysis from both time points.

### Gene expression

Total RNA was isolated from the flash-frozen right lung samples using the QIAshredder (Qiagen, cat no. 79656) and RNeasy Mini Kit (Qiagen, cat no. 74136). The tissue was weighed and homogenized using a mortar and pestle. Further steps were performed as per the instructions provided by the manufacturer. The quality and concentration of RNA were measured using the Nanodrop Spectrophotometer (Thermo Fisher Scientific, MA). Complementary DNA (cDNA) was synthesized from 170 ng of RNA using the AzuraQuant cDNA Synthesis Kit (Azura Genomics, cat no. AZ-1996) according to the manufacturer’s protocol.

Gene expression was measured for specific genes to assess senescence, fibrosis, and senescence-associated secretory phenotype (SASP) in the samples. Quantitative Polymerase Chain Reaction (qPCR) was performed using the Taqman probes (Spp1: Mm00436767_m1, Igf1: Mm00439560_m1, and IL-6: Mm00446190_m1). Tissue from sham-treated mice was pooled from both time points to increase statistical power. Rpl13a (Mm05910660_g1) was used as a housekeeping gene. The expression was calculated relative to the sham group. Data was quantified using the 2^-ΔΔCt^ method.

### Bulk RNA Sequencing

Flash-frozen right lung samples (2 samples for each condition (bleomycin- and sham-treated) and time point (D24 and D72); total n = 8) were shipped on dry ice to Azenta, USA, for RNA isolation, library preparation, and sequencing. Libraries were prepared using poly(A) selection and sequenced on Illumina NovaX-25B to generate paired-end reads. Raw reads were processed and aligned to the mouse reference genome (GRCm39) using the provider’s standard bioinformatics pipeline. Samples from sham-treated mice were pooled from both time points to increase statistical power. Differential expression analysis was performed using DESeq2, with genes considered significantly different at an adjusted p-value < 0.05 and a Log2 fold change of 1.

### Immunofluorescence Imaging

IMR-90 human lung fibroblasts (ATCC, USA; Cat# CCL-186) were cultured on collagen (Sigma-Aldrich, cat no. C0130) coated coverslips at using 5% CO_2_ and 3% O_2_ environment using Dulbecco’s Modified Eagle’s Medium (DMEM) (Corning, cat no. 10-013-CV) with 10% FBS. Senescence was induced using 300 nM of Doxorubicin (Millipore Sigma, cat no. 504042) in PD 25-35 fibroblasts for 24 hours in DMEM complete media. SEN (senescent cells) were used between 7-10 days after senescence induction. NS (non-senescent) controls were plated on day 8. SA-β-galactosidase activity was confirmed using a commercial kit (Abcam, cat. no. ab65351) as described earlier (Dimri et al., 1995). Immunofluorescence staining was done by fixing cells with 4% PFA at room temperature (RT) for 15 minutes (Thermo Scientific, cat no. AAJ19943K2). Cells were permeabilized with 0.2% Triton for 15 minutes. The samples were blocked with 5% goat serum for 1 hour at RT to reduce nonspecific binding. Rabbit anti-Spp1 primary antibody (Proteintech, cat no. 22952-1-AP) was used at a 1:300 dilution overnight at 4°C. Goat anti-rabbit Alexa Fluor 568 IgG (Invitrogen, cat no. A11036) secondary was used at 1:1000 dilution, followed by Hoechst 33342 (Invitrogen, cat no. H3570) at 1:2000 dilution for 1 hour at RT in the dark. Images were acquired on the EVOS M5000 microscope (Invitrogen) using 20x magnification. Three fields of view were obtained for each sample, and fluorescence was quantified using ImageJ. The experiment was conducted using a single biological sample (N=1).

### Statistical analysis

Statistical analysis was performed using Prism GraphPad 10 software, and significance was determined by an unpaired t-test between groups, with p values ≤ 0.05 considered statistically significant. All data are represented as the mean ± standard error of the mean (SEM) unless otherwise specified. The raw RNA-seq data have been deposited under the accession GSE####

## Results

### A Single Dose of Bleomycin Impacts Weight and Behavior in Old Mice For 72 Days

Bleomycin, a chemotherapeutic antibiotic, is an effective inducer of pulmonary fibrosis in mice and other animals due to the absence of the bleomycin-inactivating enzyme bleomycin hydrolase in the lungs (Liu et al., 2017; Rudders & Hensley, 1973). It has been previously reported that oropharyngeal administration of bleomycin in 7- to 9-week-old mice effectively induces pulmonary fibrosis in various mouse strains (Egger et al., 2013). In our previous studies, we demonstrated the successful induction of IPF in 6-month-old C57BL/6J mice using the same dosing strategy (Meca-Laguna et al., 2025a, 2025b; Barkovskaya et al., 2025). However, this model fails to capture many salient aspects of human IPF, including its chronic and progressive nature, as well as the broader effects of aging on the patient. In this study, we investigated whether a single dose of bleomycin in 17-month-old UM-HET3 mice would induce a more persistent IPF disease phenotype, thus generating a higher-fidelity model of the human disease. We decided to test two bleomycin doses: 2.5 mg/kg and 3.5 mg/kg, as previous reports suggest a dose range of 1-5 mg/kg in various mouse strains for successful IPF induction (Mehdizadeh et al., 2022; Kim et al., 2010).

A total of 37 male mice were used in the study. Animals were divided into two groups: a sham-treated group (n=9) and a bleomycin-treated group 2.5 mg/kg (n=11) or 3.5 mg/kg (n=17). Weights, body score, and spontaneous movement in the open field test (behavior) were monitored throughout the study. Mouse lungs were harvested at two time points: D24 from group 1 and D72 from group 2, to study the effect of a single dose of bleomycin on old mice, as shown in the schematic (**Figure 1A**). Mice that received bleomycin have been pooled for analysis (unless specified), as they showed similar outcomes in terms of weight loss, behavior, and overall fibrosis. Consistent with prior reports (Yang et al., 2023), our results show that 8 out of 28 mice that received bleomycin either died or had to be euthanized based on body score and weight loss in accordance with humane endpoints as per the IACUC protocol, whereas all sham-treated mice survived to the end of the study (**Figure 1B**). Bleomycin-induced IPF has been strongly correlated with weight loss and fibrotic injury in mouse models (Lamichhane et al., 2022). Consistent with this, we observed that mice treated with bleomycin lost more body weight compared to the sham-treated mice on day 24 (**Figure 1C**) and continued to lose weight until D30-35, then stabilized (**Figure 1C-i**).

**Figure 1:**
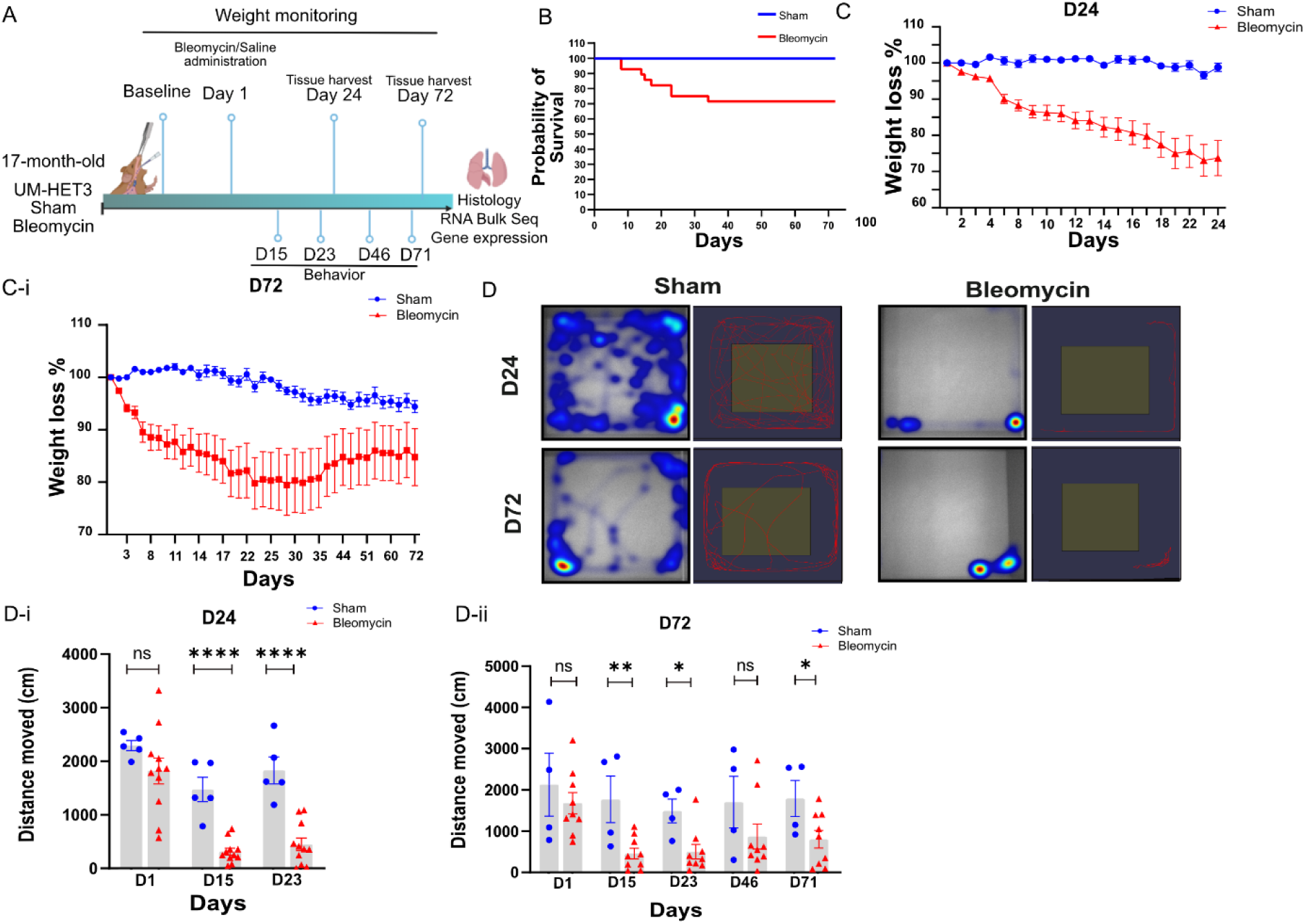
Effects of Bleomycin on Weight and Behavior in Old Mice. **(A)** Schematic representing the experimental design of fibrosis induction using a single dose of bleomycin in old male (17-month-old) UM-HET3 mice. Bleomycin was administered using the oropharyngeal route of administration. Created using BioRender.com. **(B)** Kaplan–Meier survival curves of mice treated with bleomycin. Survival was monitored over 72 days post administration. Deaths from bleomycin occurred on days 8,14,15,17,23 and 34 after bleomycin administration (death/total= 8/28), all sham mice survived till the end of study. **(C)** Bleomycin mediated body weight loss at D24. Weight is normalized to the day of bleomycin instillation (day 0), n = 16 (n = 5 sham mice; n = 11 bleomycin-treated mice). **(C-i)** Bleomycin mediated body weight loss at D72. Weight is normalized to the day of bleomycin instillation (day 1), n = 13 (n = 4 sham mice; n = 9 bleomycin-treated mice). **(D)** Heatmaps and tracking showing movement of mice in the open field arena during the 10-minute interval at D24 and D72. **(D-i)** Mouse behavior data collected at different time points (Baseline, D14, D23) recording spontaneous movement with open field for a duration of 10 minutes (n = 5 sham mice; n = 11 bleomycin-treated mice). Data is represented as mean ± SEM, unpaired t-test, ****p ≤ 0.0001. **(D-ii)** Mouse behavior data collected at different time points (Baseline, D14, D23, D46, D71) recording spontaneous movement with open field for a duration of 10 minutes (n = 4 sham; n = 9 bleomycin-treated mice). Data represented as mean ± SEM, unpaired t-test, *p ≤ 0.05, ns ≥ 0.05.

After administering bleomycin, we measured the spontaneous movement of the mice in an open field arena, a method often used as a preclinical analog for the reduced mobility of IPF patients (Shingai et al., 2021; Nishiyama et al., 2018). Our results show that mice treated with bleomycin moved significantly less compared to sham-treated mice for group 1 (D24, with a fold change of 4.4) (**Figures 1D and 1D-i**). Similarly, as shown in **Figure 1D-ii**, bleomycin-treated mice harvested at a late time point (D72, group 2) also exhibited reduced movement compared to sham-treated mice (fold change of 2.2).

### Persistent Fibrosis and Senescence in Bleomycin-Treated UM-HET3 Mouse Lungs

The lung tissue from the mice was harvested to characterize the burden of fibrosis and senescence on days 24 (group 1) and 72 (group 2) after bleomycin treatment. We employed Picrosirius red staining, a well-established histological method for assessing collagen accumulation in IPF mouse models (Vogel et al., 2015), to detect changes in matrix organization and fibrosis. **Figure 2A** confirms the presence of collagen-rich fibrotic areas in the lungs of bleomycin-treated mice, which are absent in sham-treated mice. The stained sections were then blind-scored on a scale of 1-5, ranging from low to severe fibrosis. **Figure 2A-i** represents the comparison between lung sections taken from sham- and bleomycin-treated mice at D24 (group 1), showing significantly higher fibrosis (fold change of 3.4) in bleomycin-treated lungs. We observed consistent results at D72 (group 2), with samples taken from bleomycin-treated mice demonstrating significantly higher fibrosis (fold change of 2.6) than samples from sham-treated mice (**Figure 2A-ii**). Additionally, Hematoxylin and Eosin (H&E) staining was performed to assess histological changes in the lung tissue. At both D24 and D72, tissue from bleomycin- but not sham-treated mice revealed distinct histopathological changes characteristic of IPF (**Figure 2C)**.

**Figure 2:**
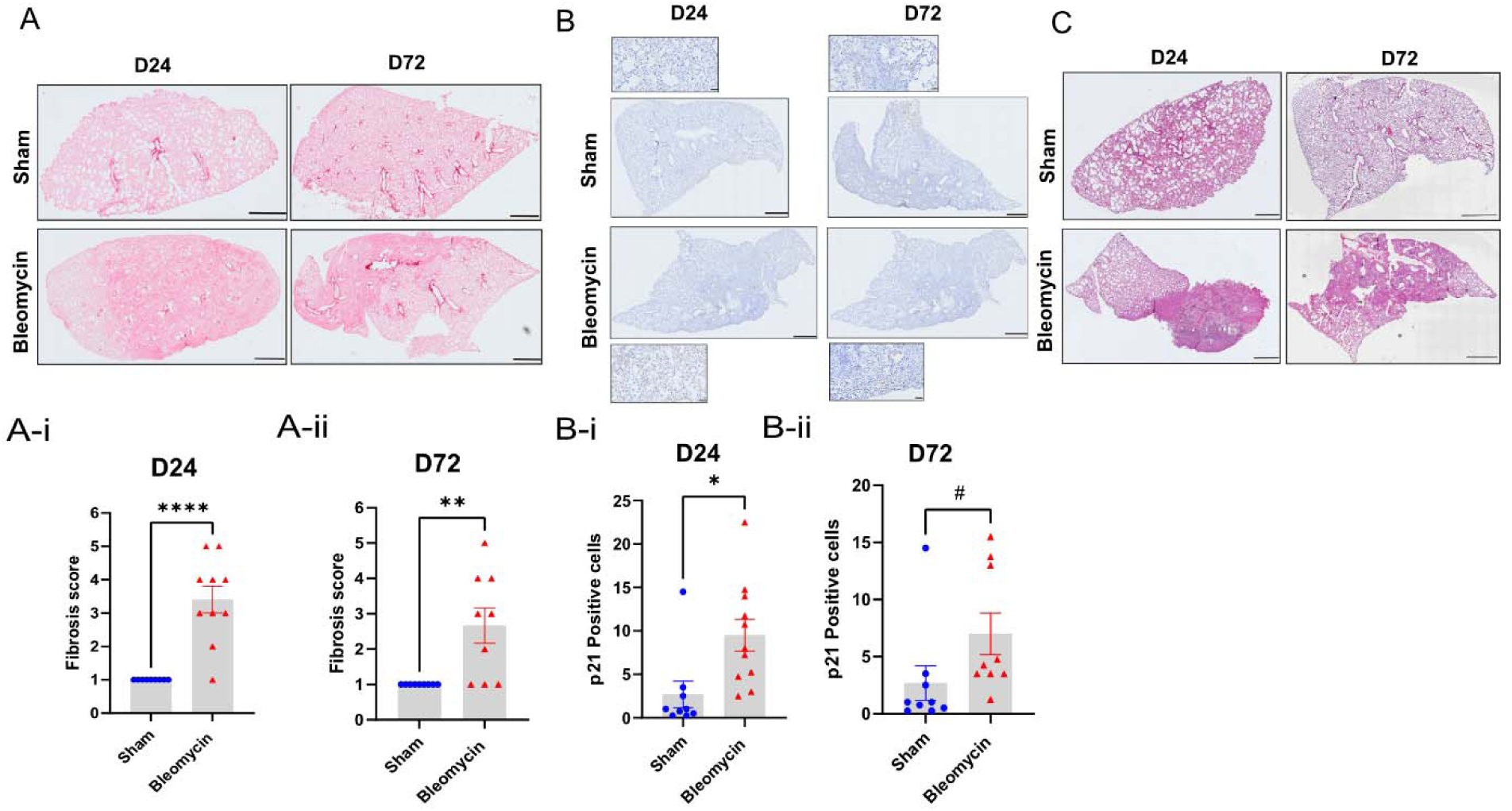
Bleomycin Treated Mice Continue to Demonstrate Injury, Fibrosis and Senescence at D72. **(A)** Representative lung sections from FFPE blocks (5 µm) stained with Picrosirius red. Scale bar = 1 mm. **(A-i)** Quantification of fibrosis in controls and bleomycin-treated mice at D24 (n = 5 sham mice; n = 11 bleomycin-treated mice) and **(A-ii)** D72 (n = 4 sham mice; n = 9 bleomycin-treated mice). Data represented as mean ± SEM, unpaired t-test. **(B)** Representative images of sham and bleomycin-treated mice IHC staining for p21^WAF1/Cip1^. Scale bar = 50 μm. **(B-i)** Quantification of p21^+^ nuclei in the left lung sections (5 µm) at D24, n = 5 saline-treated mice; n = 11 bleomycin-treated mice and **(B-ii)** D72 (n = 4 sham mice; n = 9 bleomycin-treated mice). Data represented as mean of four different fields of view per mouse ± SEM, unpaired t-test, *p ≤ 0.05, #p ≤ 0.08, ns ≥ 0.05. **(C)** Representative images of H&E-stained left lungs demonstrating bleomycin induced pulmonary damage compared to sham both days 24 and 72. Scale bar = 1 mm.

There is substantial evidence that cellular senescence plays a role in the progression of IPF (Zhang & Wang., 2023; Collard HR., 2010; Schafer et al., 2017). To assess the senescence burden in our bleomycin-treated animals, we stained sections of left lung tissue at both time points with an antibody against Cdkn1a/p21, a widely accepted biomarker of cellular senescence both *in vitro* and due to its role in cell cycle arrest and ability to integrate stress signals (Barkovskaya et al., 2025; Meca-Laguna et al., 2025a, 2025b; Wagner et al., 2022) **(Figure 2B**). IHC staining revealed a prominent increase in p21-positive cells in bleomycin-treated vs. sham-treated mice at D24 (**Figure 2B-i**) and D72 (**Figure 2B-ii**), confirming a higher senescence burden. In summary, we observed persistent weight loss, lack of spontaneous movement and lung tissue fibrosis with a senescence burden in bleomycin-treated animals, in samples collected at both the day 24 and the 72 time points, suggesting stable tissue remodeling and related deficits.

### Bulk RNA-Seq Confirms Distinct Differences Between the Two Groups

To evaluate the transcriptome of aged mice subjected to bleomycin treatment, we performed bulk RNA Sequencing (RNA-Seq) of the lungs of sham- and bleomycin-treated samples 24 and 72 days after bleomycin instillation. Principal Component Analysis (PCA) was performed to investigate clustering of samples. The PCA plot revealed a clear separation between the sham-treated and bleomycin-treated groups along the first two principal components, PC1 and PC2, that accounted for 63% and 12% of data variability, respectively (**Figure 3A**). Interestingly, samples collected at different time points occupy distinct spaces on the plot. We observe that the gene signature of the low-dose (2.5 mg/kg) bleomycin-treated mice at D72 resembles that of the sham-treated mice, suggesting substantial resolution of IPF, despite significant underlying differences in the profile between conditions at D24. Overall, PCA confirmed a considerable difference in gene expression between sham and bleomycin-treated mice at both time points. In summary, bleomycin treatment drives distinct and measurable shifts in gene expression.

**Figure 3:**
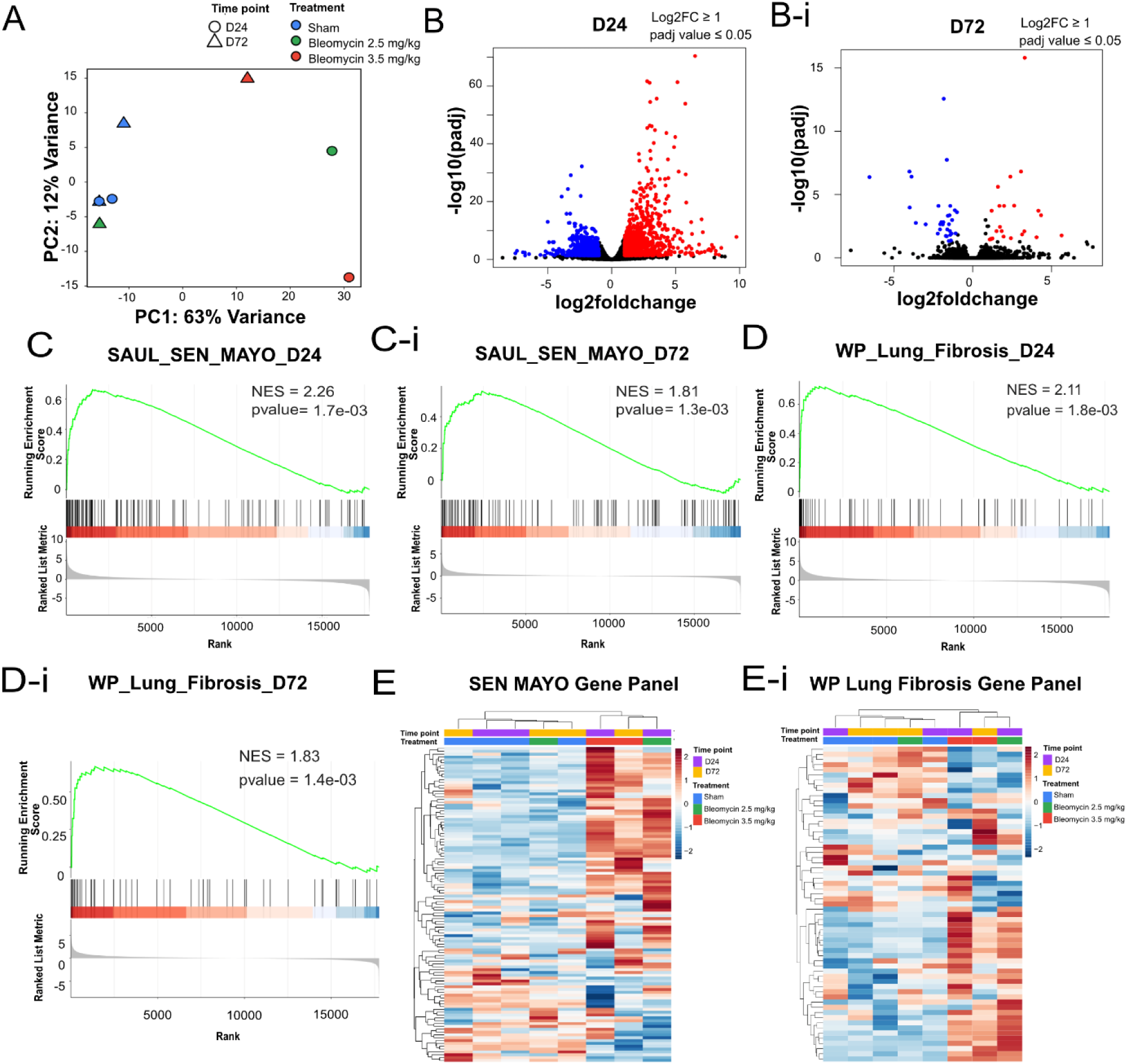
Transcriptomic Analysis Reveal Enrichment of Senescence- and Fibrosis-Associated Genes. **(A)** PCA was performed on sham (n = 2) and bleomycin (n = 2) for each time point. Colors indicate different doses of bleomycin and shapes denote the time point. **(B)** Volcano plot showing differentially expressed genes between sham and bleomycin samples at D24 and **(B-i)** D72. Red for upregulated genes and blue for downregulated genes. **(C, C-i)** GSEA of sham mice compared to bleomycin treated mice for the “SAUL SEN MAYO” gene set at D24 and D72 shows senescence signature in fibrotic samples and **(D, D-i)** GSEA of sham mice compared to bleomycin treated mice of “WP LUNG FIBROSIS” gene set at D24 and D72. **(E)** Heat map displaying expression profiles of genes from the SEN MAYO senescence signature across sham and bleomycin-treated at D24 and D72. **(E-i)** Heatmap of expression profiles of genes between sham and bleomycin-treated at D24 and D72 using the WP Lung fibrosis gene set.

Differentially expressed genes (DEG) were visualized using a volcano plot at D24 (**Figure 3B**) to highlight genes that were significantly altered between the bleomycin- and sham-treated lung tissue samples. A total of 1652 genes were identified at D24, among which 941 genes were significantly upregulated, and 711 genes were downregulated in the bleomycin group. A smaller number of DEGs (54 total; 23 upregulated and 31 downregulated) were seen at D72 (**Figure 3B-i**). However, for D72, a high dose of bleomycin (3.5 mg/kg) resulted in 2,122 DEGs being observed. In addition, gene set enrichment analysis (GSEA) of lungs from sham- and bleomycin-treated mice demonstrated enrichment of multiple genes related to senescence and fibrosis.

To compare our findings with a well-characterized senescence gene signature, we utilized the SenMayo gene set, derived from Mayo Clinic transcriptomic data (Saul et al., 2022). The genes differentially expressed in bleomycin (both doses)- vs. sham-treated lungs exhibited a strong enrichment signal, indicated by a higher normalized enrichment score (NES) of 2.26 (**Figure 3C**) and 1.81 (**Figure 3C-i**) at days 24 and 72, respectively. These findings further support the presence of a senescence phenotype in fibrotic lung tissue, consistent with the known role of cellular senescence in the pathogenesis of IPF.

We also compared the lung tissues of sham and bleomycin-treated mice (at both doses) to investigate the activation of fibrosis-related pathways. The WP_LUNG_FIBROSIS gene set revealed a clear enrichment of fibrotic signaling at D24 and D72 with an NES of 2.11 and 1.83, respectively (**Figures 3D, 3D-i**). This indicates robust activation of genes involved in extracellular matrix remodeling, myofibroblast activation, and pro-fibrotic cytokine signaling at D24 (both doses) and D72 (high dose only) (**Supplementary Figures 1A, 1A-i, 1A-ii and 1B, 1B-i, 1B-ii**).

To further investigate transcriptional changes in the bleomycin- compared to the sham-treated samples, we performed gene expression analysis across the samples. A heat map of 100+ genes from the SenMayo panel is shown in **Figure 3E**. The groups are categorized based on treatment (sham vs. bleomycin), dose (2.5 mg/kg vs. 3.5 mg/kg), and time point (D24 vs. D72). A clear distinction was seen between the sham- and bleomycin-treated samples at D24. At D72, the mice treated with 3.5 mg/kg of bleomycin noticeably grouped differently from the sham-treated mice, whereas the mice treated with 2.5 mg/kg grouped more closely with the sham-treated mice. This indicates a visible and distinct response to treatment, and partial resolution over time in the low-dose bleomycin group, consistent with our pathological findings. We also explored a similar analysis of the genes in the WP_LUNG_FIBROSIS gene set, and similar patterns were observed between the sham- and bleomycin-treated samples (**Figure 3E-i**).

### Bleomycin-Induced IPF UM-HET3 Mice Share Common Senescence Genes with Other Mouse Strains and Human Idiopathic Pulmonary Fibrosis (IPF)

Our objective was to compare the gene expression signature of late-life induced model IPF UM-HET3 mice with that of different mouse strains and human IPF to identify conserved disease pathways and genes. We searched for SenMayo genes in our differentially expressed genes (DEGs) at D24 and generated a list of genes that are highly expressed in the bleomycin-treated samples (**Figure 4A**). Similarly, the leading-edge genes in the 3.5 mg/kg bleomycin-treated samples were analyzed using the SenMayo gene set to determine a list of genes that are highly expressed at D72 (**Figure 4A-i**). A Venn diagram comparing differentially expressed genes (DEGs) in our D24 mice with those in other experimental fibrosis models using C57BL/6 mice at D21, as re-analyzed from Peng et al. (2013), is presented in **Figure 4B**. The diagram illustrates the overlap of DEGs from the SenMayo set identified by RNA-Seq in both models (Spp1, Ereg, Igf1, Il6, Ccl8, Mmp12, Mmp13, Cxcl10, Mmp2, Cxcl1, Ccl3). A similar analysis of the overlap of DEGs from human IPF tissue (Schafer et al., 2017), the SenMayo gene panel, and our UM-HET3 data from D24 and D72 (high dose, 3.5 mg/kg) is represented in **Figure 4B-i**. This confirms an overlap of some genes between our aged IPF model and genes broadly differentially expressed in senescent cells across various cell types of origin and modes of induction, as well as in human IPF. This comparative analysis supports the translational relevance of our aged IPF model and could assist in identifying targets potentially involved in IPF pathogenesis.

**Figure 4:**
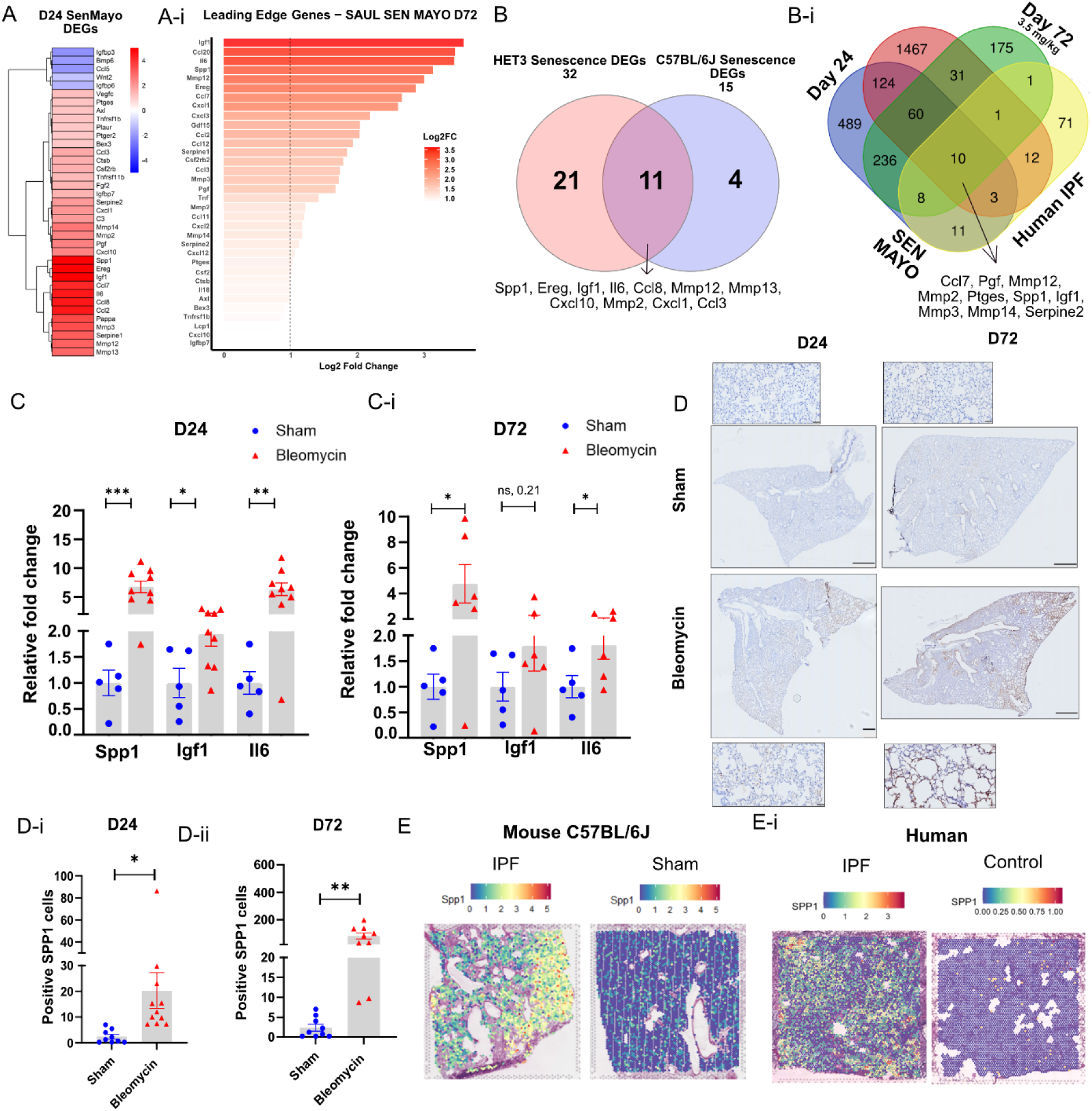
Gene Expression and Spatial Transcriptomics Confirm Higher Expression of SEN Mayo Genes in Bleomycin Treated Samples. **(A)** Senescent genes identified among the DEGs at D24 from the SEN MAYO panel. **(A-i)** Leading edge genes identified in the SEN MAYO panel at D72. **(B)** Venn diagram indicating an overlap of DEGs between C57BL/6J and HET3 mice using the SEN MAYO senescence gene set. 11 genes from the SEN MAYO panel were found to be present in both sets. C57BL/6J data re-analyzed from Peng et al., 2013. **(B-i)** Venn diagram comparing genes from Bleomycin-treated mice at D24 and D72 (3.5 mg/kg) with SEN MAYO senescence gene panel and Human IPF data. Data was re-analyzed from Schafer et al., 2017. **(C, C-i)** Gene expression of Spp1, Igf1 and Il6 of sham and bleomycin-treated mice. Rpl13a was used as housekeeping control. Significance calculated using unpaired t-test; data represented as mean ± SEM at D24 and D72. **(D)** Representative IHC images at D24 and D72 of sham and bleomycin-treated mice stained for SPP1. Scale bar = 50 um. **(D-i)** Spp1 positive cell quantification at D24 and **(D-ii)** D72. Data represented as mean of four different fields of view per mouse ± SEM, unpaired t-test, *p ≤ 0.05, ** p ≤ 0.002, ns ≥ 0.05. **(E)** Validation of Spp1 expression data via spatial transcriptomic analysis in mouse IPF samples. Mouse samples with IPF showcase higher expression of Spp1. **(E-i)** Validation of SPP1 expression in human IPF samples. Data was re-analyzed from Franzen et al., 2024 for both human and mouse tissues. IPF = Freshly frozen tissue. IPF lung tissue collected with severe tissue remodeling. Control = Healthy lung tissue no known evidence of lung disease.

A subset of genes from the UM-HET3 SenMayo Leading Edge, specifically those genes contributing significantly to the enrichment score, was selected for validation by RT-qPCR. As illustrated in **Figure 4C**, the genes Spp1, Igf1, and Il-6 were significantly upregulated in bleomycin- compared to sham-treated samples. At D72, all genes remained upregulated in bleomycin-treated samples, consistent with earlier findings (**Figure 4C-i**). We further validated SPP1 expression using IHC, showing a higher proportion of SPP1-positive cells in bleomycin-treated samples compared to sham-treated samples (**Figure 4D).** The quantitative analysis further confirmed that a significantly higher percentage of SPP1-positive cells were present in bleomycin-treated samples at D24 and D72 (**Figures 4D-i, D-ii**). Re-analysis of human and mouse spatial transcriptomic data confirmed the presence of SPP1 spots (**Figure 4E**) (Franzén et al., 2024). Additionally, SEN and NS controls were stained for SA-β-gal *in vitro* (**Supplementary Figure 2A, A-i**), and we found higher SPP1 levels in SEN compared to NS controls via immunofluorescence (**Supplementary Figure 2B, B-i**). In summary, we report the use of the old UM-HET3 IPF mouse model as a clinically relevant model that better mimics human IPF than the standard bleomycin-induced mouse model.

## Discussion

IPF is a fatal lung disease that mainly affects the aged population and has limited medical options. However, the bleomycin-induced senescence model primarily represents an acute injury with a repair process (Walters et al., 2008; Chung et al., 2003), which makes it a poor model for the chronic, progressive, and heterogeneous nature of human IPF.

Different dosing strategies and administration routes, such as intraperitoneal injection, intratracheal administration, intravenous, intranasal, and oropharyngeal aspiration, have been evaluated in young mice (Gul et al., 2023; Egger et al., 2013). However, most studies use mice aged 6–8 weeks, which are developmentally analogous to adolescent or young adult humans, significantly younger than the typical age of IPF onset (Jenkins et al., 2017). Further efforts to refine the model have involved administering bleomycin to young and old mice in multiple doses or repeated cycles to promote more progressive and chronic fibrosis. However, this strategy is more time-consuming and does not capture the effects of biological aging on human IPF patients, which profoundly impact their clinical condition (Redente et al., 2021; Collard HR., 2010; Degryse et al., 2010). Additionally, to the best of our knowledge, all previous bleomycin-induced mouse models have been implemented in established mouse strains, which are genetically identical, making the generalizability of findings questionable even among mouse strains (Phan & Kunkel, 1992), let alone their translatability to genetically diverse human populations (Jenkins et al., 2017; Moore et al., 2008).

The lack of a robust IPF model that mimics human pathology hinders clinical development. To address these limitations, we developed an improved aged mouse model of IPF in which a single dose of bleomycin is administered to 17-month-old UM-HET3 mice. In contrast to conventional bleomycin mouse models, which are known to self-resolve (Li et al., 2023; Zia et al., 1992), this approach more closely mimics the chronic and persistent nature of human IPF. This approach distinguishes our study from previous studies that either rely on specific strains of mice with variable susceptibility to bleomycin (Walkin et al., 2013; Phan & Kunkel, 1992) or employ cumbersome repeat-dose strategies.

In this study, we investigated physiological, behavioral and molecular alterations at two distinct timepoints: an early stage (Day 24 post-bleomycin administration), by which point fibrosis typically begins to resolve in conventional bleomycin models spontaneously (Song et al., 2022; Kolb et al., 2020); and a late stage (Day 72), to confirm the persistence of IPF-associated pathology. Our study confirms that bleomycin-induced pulmonary fibrosis results in significant weight loss in mice at both early and late stages. This aligns with prior reports indicating that weight loss is a systemic effect of bleomycin exposure. Mice do not exhibit a complete recovery of weight, which likely reflects a combination of systemic inflammation, reduced food intake, and increased catabolic activity linked with disease progression (Manali et al., 2011; Inayama et al., 2006). Furthermore, the persistence of fibrosis has been confirmed by staining collagen in the lung tissue. We observed that high degrees of fibrosis persist in bleomycin-treated aged mice through at least D72. Finally, we evaluated the spontaneous activity and fatigue using the open field test, which is reduced because of lung dysfunction in bleomycin-treated mice, as in humans with IPF (Kahlmann et al., 2020)

Previous studies have reported reduced overall movement in young (4–6-month-old) C57BL/6J mice treated with bleomycin (Shafer et al., 2017; Meca-Laguna et al., 2025a, 2025b). In this study, we evaluate distance traveled in the open field at different time points throughout the study as a measure of lung injury and systemic inflammation, using it as an effective means to track the progression of the disease over time. We report a reduction in spontaneous activity following bleomycin induction. This loss of activity was apparent even at D72, suggesting a persistent and non-resolving IPF phenotype.

It is well documented that senescent cells accumulate in different tissues with age and afflicted tissues in multiple age-related diseases (Tuttle et al., 2021; Muñoz-Espín & Serrano., 2014) and have been specifically implicated in the pathogenesis of IPF (Zhang & Wang., 2023; Yao et al., 2021; Schafer et al., 2017). The role of senescent cells in the progression of fibrosis is well-established, although their role in the resolution of injury is complex (Redgrave et al., 2023; Meyer et al., 2016; Demaria et al., 2014; Krizhanovsky et al., 2008), and there is limited understanding of their specific role in IPF. Pulmonary AT2 cells, endothelial cells, and fibroblasts have been reported to undergo senescence in the lungs of individuals with IPF (Liu and Liu, 2020). Pre-clinical studies have shown that selectively ablating senescent cells can improve lung and physical function in model IPF, whether via senolytic drugs (such as navitoclax, ABT-263 — Pan et al., 2017), the cocktail of dasatinib plus quercetin (D+Q — Schafer et al., 2017), or the PIKfyve inhibitor apilimod (Barkovskaya et al., 2025) or via adoptive immunosurveillance with γ δT cells (Meca-Laguna et al., 2025b). A pilot clinical study suggested that D+Q improved physical function in human IPF patients (Justice et al., 2019), although a small phase I trial did not detect a significant difference (Nambiar et al., 2023).

In the present study, lung tissue from bleomycin-treated mice exhibited an increase in p21-positive cells upon IHC staining, confirming the presence of senescent cells in the IPF model after 72 days of treatment. Additionally, we observed a clear enrichment of senescence-associated gene expression pathways in the aged IPF UM-HET3 mice, which were differentially expressed compared to the baseline expression of these pathways in the aged sham-treated controls. We shortlisted conserved senescence-associated genes between human IPF patients and bleomycin-induced UM-HET3 and C57BL/6J model IPF.

Our RNA-Seq transcriptomic results indicate an increased expression of senescence- and fibrosis-associated genes in bleomycin-treated mice compared to sham-treated mice. PCA and heat maps confirm that bleomycin-treated mice cluster differently and have different gene expression profiles compared to sham-treated mice. However, the samples taken from mice treated with 2.5 mg/kg of bleomycin appear to cluster closer to the sham-treated mice, which could mean that a higher dose of 3.5 mg/kg may be more effective in inducing persistent fibrosis in mice.

Furthermore, sham- and bleomycin-treated tissue harvested on days 24 and 72 revealed a set of genes that are highly expressed in our samples, which are present in the SenMayo panel. We report elevated expressions of Spp1, Igf1, Mmp2, Ccl2, Ccl3, and Il6 in our D24 samples and D72 high-dose (3.5 mg/kg) samples. Notably, these genes were also upregulated in our previous study using 6-month-old C57BL/6J mice (Meca-Laguna et al., 2025a). Gene expression analysis of these genes revealed significant upregulation at both time points, corroborating the RNA-Seq findings. Osteopontin or Secreted Phosphoprotein 1 (SPP1) promotes cell proliferation and inhibits apoptosis through NF-kB activation (Standal et al., 2004). In the context of IPF, SPP1 is believed to play a role in fibroblast proliferation, aggregation, and accumulation via the PI3K/Akt/mTOR pathway, and elevated levels of SPP1 have been observed in IPF patients and mouse samples (Yue et al., 2024). Macrophages with high SPP1 expression are found to be abundant in human IPF samples, which could be involved in the progression of the disease (Morse et al., 2019). In a recent study, SPP1 was identified (among others) as a vascular specific senescent cell marker based on transcriptional profiles (via the SenMayo GSEA) in a murine atherosclerotic remodeling. These cells failed to express traditional markers of senescence such as p16 or p21. Treatment with a senolytic, ABT-737 showed a decrease in SPP1 positive senescent cells suggesting its potential as a therapeutic target (Mazan-Mamczarz et al., 2025).

Similarly, Insulin-like Growth Factor-1 (IGF1) has been reported to be involved in cell proliferation and activation of fibroblasts to the myofibroblast phenotype. It is believed to lead to the senescence of alveolar epithelial cells (AECs), thereby directly aiding disease progression and indirectly via paracrine effects on fibroblasts (Sun et al., 2021). Our data also confirms the increased expression of Igf1 in bleomycin-treated mice.

Additionally, matrix metallopeptidases (MMPs), a family of zinc-dependent endopeptidases, have been extensively studied for their role in regulating extracellular matrix (ECM) remodeling. In addition to its classical role, Mmp2 has been linked to aberrant activation of the Wnt/β-catenin signaling pathway, a process implicated in IPF (Bormann et al., 2022; Craig et al., 2015).

Our RNA-Seq data also showed high levels of chemokines. Numerous chemokines have been implicated in IPF in animal models and prospective studies via their roles in promoting inflammation, leukocyte recruitment, and fibrogenic signaling through receptor-mediated activation of lung and immune cells (Russo & Ryffel, 2024). CCL3 (MIP-1α), one of the chemokines implicated in our model and overlapping with the SenMayo panel, is a proinflammatory chemokine responsible for recruiting immune cells, such as CD8⁺ T cells, neutrophils, and macrophages, through its interaction with CCR5. Antibody-mediated blockade of CCL3, genetic deletion of CCL3, or genetic deletion of CCR5 mitigates fibrosis and leukocyte infiltration in mouse IPF models, highlighting CCL3 as a key mediator in IPF pathogenesis and a promising therapeutic target (Liu et al., 2023). Studies have reported the involvement of another of the DEGs identified in our IPF model mice, CCL2, in enhancing TGF-β1 expression in pulmonary fibroblasts and indirectly promoting collagen synthesis in isolated rat lung fibroblasts through autocrine or juxtacrine signaling loops (Gharaee-Kermani et al., 1996).

IL-6, the last of our top DEGs that are included in the SenMayo panel, has also been suggested to play a causal role in IPF pathogenesis. In bleomycin-treated mice, M2-like macrophages promote formation of the IL-6/sIL-6Rα complex, triggering IL-6 trans-signaling in lung fibroblasts and other cells, consequently enhancing cell proliferation and ECM production (She et al., 2021).

Our pathway analysis shows the involvement of these genes in lung fibrosis in both mice and human IPF patients (**Supplementary Figures 1C, 1C-i**). Additionally, the overlap of our DEGs with genes identified in other studies involving the C57BL/6J mouse model (**Figure 4B**) or human IPF (**Figure 4B-i**) highlights conserved molecular signatures between these models. Further exploration of these common gene signatures could yield valuable insights into the underlying mechanisms and identify novel intervention strategies.

In conclusion, this study presents a bleomycin-induced pulmonary fibrosis model in aged, genetically diverse mice at a lower cost and with less difficulty than the main alternatives that exhibits sustained fibrotic and senescent phenotypes, which better parallel those observed in human IPF while remaining inexpensive and convenient for investigators. The transcriptomic, behavioral, histological, and molecular end points shared between this new model, previous IPF mouse models, and human IPF patients provide a foundation to understand disease pathogenesis and commend this model for use in testing interventions, hence creating a promising avenue for future research.

## Acknowledgements

We thank Retro Bio, USA, for their generous donation of UM-HET3 mice. We would also like to thank Esmeralda Jimenez for her help with animal maintenance, welfare checks, and assistance with mouse experiments. Finally, we thank the donors of Lifespan Research Institute and SENS Research Foundation for their support.

## Author contribution

A. Shankar, G.M.L., and A.B. performed experiments and analyzed results. A. Sharma and A. Shankar designed experiments. A. Shankar and A. Sharma wrote the manuscript. M.J.R. provided valuable input and extensively edited the manuscript. A. Sharma obtained funding and supervised the project. All authors contributed to the discussion. All authors have read the manuscript and agree with the final version of the manuscript.

## Data availability

Data will be made available upon request.

