## Supplementary material for "The Old Genetically Heterogeneous Mouse Model Recapitulates Chronic and Persistent Idiopathic Pulmonary Fibrosis with Strong Senescence Signatures": Supplimental Figures Legends

**Supplementary Figure 1: Pathway analysis of DEGs identified from D24 and D72**

**(A, A-i, A-ii)** Wikipathways, Reactome, and Gene Ontology pathways of the 941 DEGs (out of 1652 total genes) identified at D24. **(B, B-i, B-ii)** Wikipathways, Reactome, and Gene ontology pathways of the 522 DEGs identified (1006 total genes) at D72 from samples treated with 3.5 mg/kg of bleomycin. **(C)** Mouse pathway enrichment analysis of the senescence genes identified at D72 (3.5 mg/kg). **(D)** Human pathway enrichment analysis of the senescence genes identified at D72 (3.5 mg/kg).

**Supplementary Figure 2: Validation of high SPP1 expression in senescent cells**

Senescent cells (SEN) treated with doxorubicin confirmed significantly increased SPP1 (osteopontin) expression compared to non-senescent (NS) controls. **(A)** Representative images of SA-β Gal staining showing higher activity in SEN. **(A-i)** Quantification of SA-β Gal activity confirms higher activity in SEN (N=1). **(B)** Representative images at 20x magnification of SPP1 expression in SEN and NS controls. **(B-i)** Quantification of SPP1 expression using corrected total cell fluorescence (CTCF), N=1.
